# Comparison of T1- and T2-weighted MRI contrasts of *ex vivo ex situ* brains fixed with solutions used in gross anatomy laboratories

**DOI:** 10.1101/2025.10.13.682071

**Authors:** Eve-Marie Frigon, Victoria Perreault, Amy Gérin-Lajoie, Liana Guerra Sanches, Roqaie Moqadam, Yashar Zeighami, Denis Boire, Mahsa Dadar, Josefina Maranzano

## Abstract

Post-mortem magnetic resonance imaging (MRI) offers high resolution and histological correlation, so protocols have been developed by brain banks using hemispheres fixed by immersion in Neutral-buffered formalin (NBF), but they provide limited tissue samples. Conversely, anatomy laboratories could supply complete brains perfused either with a salt-saturated (SSS) or an alcohol-formaldehyde (AFS) solution. These fixation methods alter the brain’s molecular properties, potentially affecting MRI quality and structural characteristics. T1- and T2-weighted (T1w, T2w) contrasts change with NBF fixation, but the effects of SSS or AFS remain unknown. We compared T1w and T2w intensities of different regions of interest (ROIs), including subcortical white matter (WM), cortical and deep gray matter (GM), in brains fixed with NBF, SSS and AFS.

We scanned 20 *ex situ* hemispheres (NBF-immersed=7; SSS-perfused=7; AFS-perfused=6) in a 3T MRI scanner using T1w (0.7mm^3^) and T2w (0.64mm^3^) sequences overnight. Mean intensities of 29 ROIs in T1w and T2w MRIs and GM-WM ratios were calculated and compared in brains fixed with the three solutions.

We found that T1w images were more affected by the fixation process, inverting the contrast of *in vivo* T1w and reducing the GM-WM contrast in AFS-fixed brains. T2w images resembled *in vivo* scans and maintained a sharp contrast in brains fixed with the three solutions, although the GM-WM intensity ratios were lowered in SSS-fixed brains.

In conclusion, brains fixed with SSS and AFS from anatomy laboratories could be used for MRI studies, especially with the T2w sequence that seems more appropriate for structural analyses in different ROIs.

## Introduction

Magnetic resonance imaging (MRI) techniques are widely used *in vivo* to assess the main tissue types of the brain, namely gray matter (GM) (either cortical GM [cGM] or deep GM [dGM]) and white matter (WM) [1]. MRI reveals intensity changes in different GM and WM structures during normal or pathological aging [2-4]. MRI of post-mortem brains offers advantages over *in vivo* analyses, notably its direct association with histology that remains the Gold Standard method for validation of MRI findings [5, 6]. Indeed, histology shows the cellular and molecular composition of the corresponding MRI signal in normal GM and WM regions as well as in lesions observed in neuropathology [7-9]. Furthermore, without time constraints or artifacts associated with patient movement, it is possible to obtain higher resolution and quality of the images [10-14].

For *ex vivo* studies, brain tissue is generally provided by brain banks, which separate the two hemispheres sagittally, one of which is frozen and the other immersed in neutral-buffered formalin (NBF). Brain banks usually provide researchers with few and small tissue samples from the chemically fixed hemisphere, limiting MRI studies that require full brains. As such, anatomy laboratories could become a potential source of full brains for neuroscientists, but these are fixed by whole-body perfusion with solutions commonly used for gross-anatomy labs, such as a salt-saturated solution (SSS) or an alcohol-formaldehyde solution (AFS). In perfusion fixation, the fixative diffuses from the vessels, which are distributed throughout the brain parenchyma, resulting in distinct patterns of fixation, while in immersion, the fixative diffuses only from the surface to the center of the brain [15]. This may produce a poorer fixation of deep cerebral structures, such as dGM nuclei, and over-fixation of superficial regions such as cGM and subcortical WM (scWM) [15, 16]. In addition, SSS and AFS have effects on the cellular and molecular properties of brain tissue [17, 18] that could differentially affect the MRI signal.

Prior work, including ours, has made significant progress in developing MRI protocols for scanning *ex vivo ex situ* cerebral hemispheres fixed with NBF [10, 12, 14, 19], showing an impact of NBF fixation on T1- and T2-weighted (T1w, T2w) tissue contrasts. NBF changes the molecular properties of post-mortem tissue and consequently MRI intensities and relaxation times longitudinally as the fixative diffuses into the brain [20-24]. Furthermore, changes in voxel intensities (because of chemical fixation) may affect, for example, the volumetric estimation of structures measured by intensity-based segmentation protocols [27, 28]. Specifically, post-mortem NBF-fixed brains show an inverted T1w cGM-WM contrast, while T2w contrasts are not significantly affected and resemble *in vivo* scans [10, 29, 30]. Consequently, it is to be expected that brains fixed with anatomy laboratory solutions (i.e., SSS and AFS) would show similar changes of GM and WM contrasts on MRI. To the best of our knowledge, there are no studies on the impact of different fixative solutions on MRI T1w or T2w intensities and their suitability for *ex vivo ex situ* MRI studies. In this study, we examine whether brains fixed in anatomy laboratories with two different chemical solutions (i.e, SSS, or AFS) show voxel intensities comparable to those of brains fixed by immersion in NBF across multiple regions of interest (ROIs).

## Methods

### Population

We used a convenience sample of 13 *ex situ* human brain hemispheres from the body donation program of the Université du Québec à Trois-Rivières (UQTR) (SSS: N=7; AFS: N=6) that were fixed *in situ* by whole-body perfusion. We also obtained 7 specimens from the Douglas-Bell Canada Brain Bank (DBCBB), fixed *ex situ* by immersion in NBF, and matched, as closely as possible, for age, PMI, and FL. Prior to their death, donors consented to the donation of their body for teaching and research purposes, and to the sharing of their medical information. This study was approved by UQTR’s Ethic Subcommittee on Anatomical Teaching and Research and by the Douglas Research Centre Ethic Committee (McGill University, Montreal). The donor’s characteristics and mean per group are reported in Table 1.

**Table 1.**
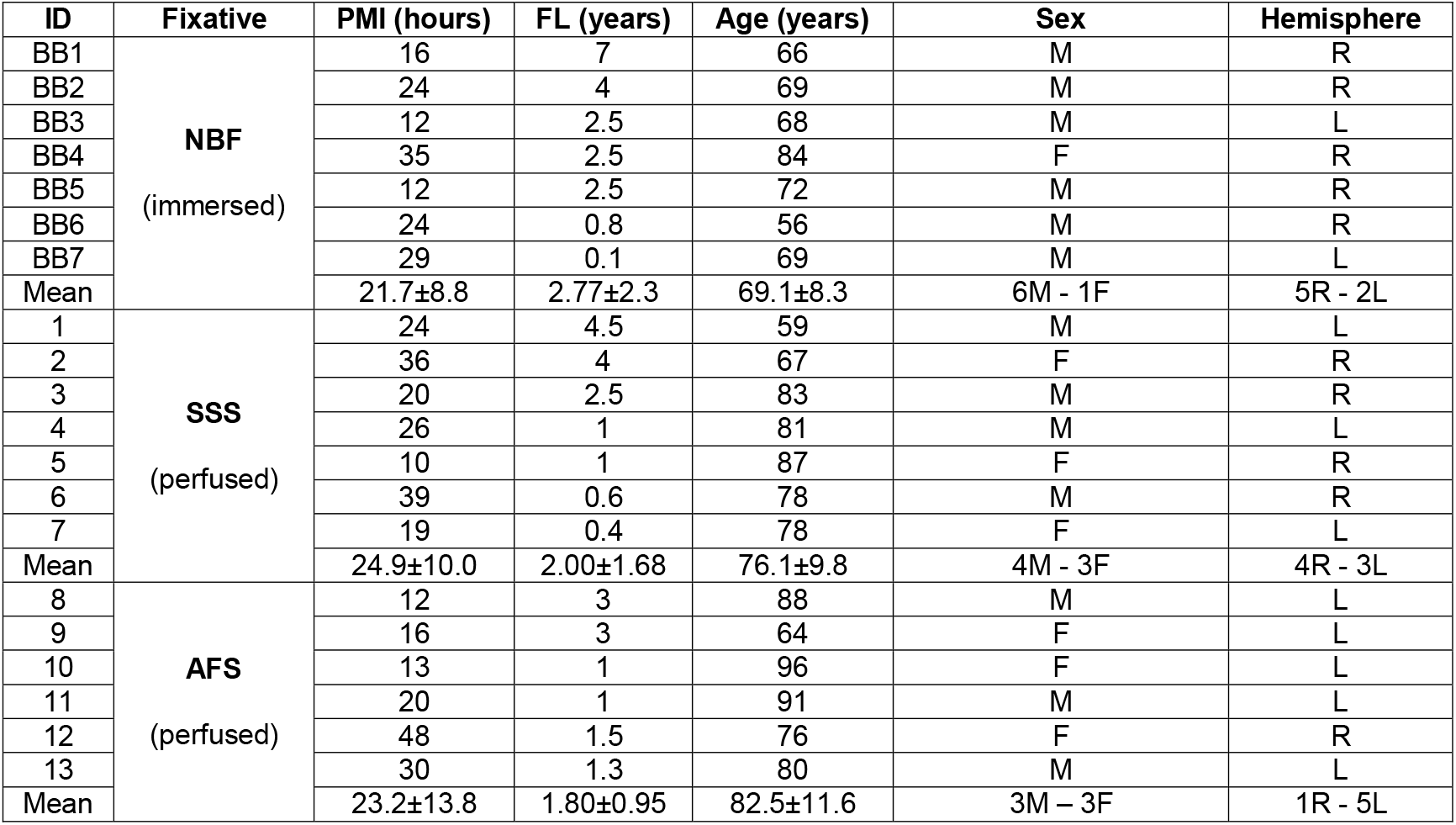
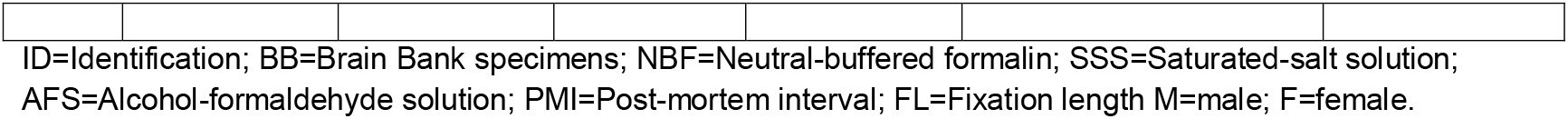
Demographic variables of the processed hemispheres.

| ID | Fixative | PMI (hours) | FL (years) | Age (years) | Sex | Hemisphere |
| --- | --- | --- | --- | --- | --- | --- |
| BB1 | NBF<br>(immersed) | 16 | 7 | 66 | M | R |
| BB2 |  | 24 | 4 | 69 | M | R |
| BB3 |  | 12 | 2.5 | 68 | M | L |
| BB4 |  | 35 | 2.5 | 84 | F | R |
| BB5 |  | 12 | 2.5 | 72 | M | R |
| BB6 |  | 24 | 0.8 | 56 | M | R |
| BB7 |  | 29 | 0.1 | 69 | M | L |
| Mean |  | 21.7±8.8 | 2.77±2.3 | 69.1±8.3 | 6M - 1F | 5R - 2L |
| 1 | SSS<br>(perfused) | 24 | 4.5 | 59 | M | L |
| 2 |  | 36 | 4 | 67 | F | R |
| 3 |  | 20 | 2.5 | 83 | M | R |
| 4 |  | 26 | 1 | 81 | M | L |
| 5 |  | 10 | 1 | 87 | F | R |
| 6 |  | 39 | 0.6 | 78 | M | R |
| 7 |  | 19 | 0.4 | 78 | F | L |
| Mean |  | 24.9±10.0 | 2.00±1.68 | 76.1±9.8 | 4M - 3F | 4R - 3L |
| 8 | AFS<br>(perfused) | 12 | 3 | 88 | M | L |
| 9 |  | 16 | 3 | 64 | F | L |
| 10 |  | 13 | 1 | 96 | F | L |
| 11 |  | 20 | 1 | 91 | M | L |
| 12 |  | 48 | 1.5 | 76 | F | R |
| 13 |  | 30 | 1.3 | 80 | M | L |
| Mean |  | 23.2±13.8 | 1.80±0.95 | 82.5±11.6 | 3M - 3F | 1R - 5L |

### Imaging protocol

The brain hemispheres were scanned using a 3T Siemens Prisma Fit human MRI scanner at the Douglas Cerebral Imaging Centre (CIC), using T1w and T2w sequences based on protocols developed at the CIC [10]. The sequences were as followed: 3D T1w MPRAGE (TR=2400ms; TE=2.41ms; FA=8deg; Resolution=0.7x0.7x0.7mm; Bandwidth=210Hz/Px; Acquisition time=7:40minutes) and 3D T2-SPACE (TR=2500ms; TE=198ms; FA=120; Resolution=0.64x0.64x0.64mm; Bandwidth=625Hz/Px; Acquisition time=7:38minutes).

### Image processing

All images were then pre-processed using the following steps: image denoising [31], intensity inhomogeneity correction [32], and image intensity normalization into range (0-100). The T1w and T2w images were co-registered and then linearly registered to the MNI-ICBM152-2009c average template [33].

### Variables of interest

T1w and T2w images were qualitatively assessed using two categorical variables: 1) Inverted cGM-WM contrast, since it was shown that post-mortem T1w images have an inverted contrast [10, 29] that may affect the accuracy of cGM and WM segmentation (Figure 1A,B) and 2) GM-WM contrast sharpness, because the degree of sharpness is also related to the accuracy of the segmentation of these tissue types (Figure 1C,D,E). These two variables were assessed in the cGM and scWM in the six different lobes (i.e., frontal, parietal, temporal, occipital, insular, cingulum) and between dGM structures (i.e., putamen, external and internal globus pallidus, claustrum, thalamus, caudate, hippocampus, amygdala) and the surrounding deep WM (dWM) (corpus callosum, fornix, internal capsule).

**Figure 1.**
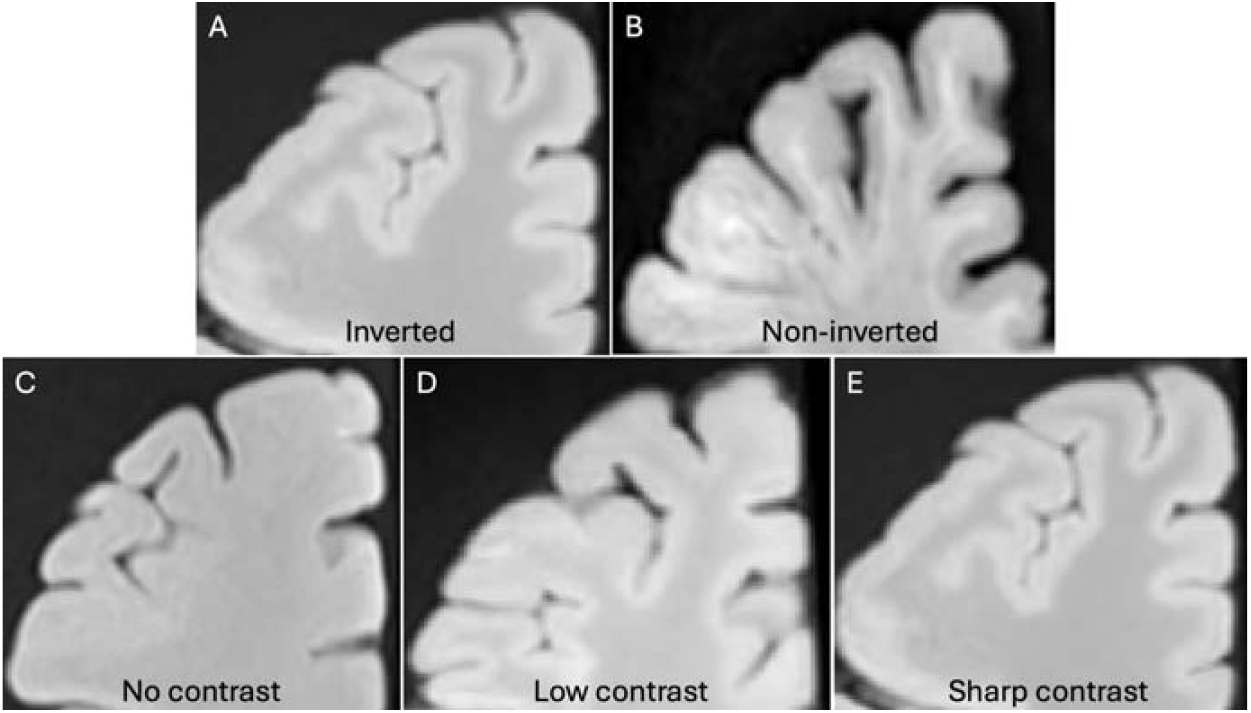
Visual assessment of the qualitative variables of interest. Variable 1 represents the Inverted cGM-WM contrast (A) or Non-inverted contrast (B). Variable 2 represents the cGM-WM contrast sharpness, where we could find no contrast (C), a low contrast (D) or a sharp contrast (E).

Variable 1: Inverted cGM-WM contrast

0=yes (Figure 1A)

1=no (Figure 1B)

Variable 2: GM-WM (cGM-WM and dGM-WM) contrast sharpness

0=no contrast (Figure 1C)

1=low contrast (Figure 1D)

2=sharp contrast (Figure 1E)

We also assessed the voxels’ intensities quantitatively by creating masks of ROIs in 29 different neuroanatomical structures, selected by a neuroanatomist with over 7 years of expertise in MRI segmentation. The ROIs were created using Display software (MINC Toolkit), by marking 500 voxels per structure to allow for accurate estimation of regional intensities. Structures selected were grouped into 1-cGM per lobe; 2-NAWM per lobe (we excluded regions that were detected as white matter hyperintensities, which are frequent in aging populations such as ours) [28]; 3-dGM; and 4-dWM. Specific regions per groups and number of voxels per masks are presented in Table 2. The voxel intensities and the mean ROI intensity values were extracted automatically from the T1w and the T2w images using MATLAB (see Supplementary Tables 1-2 for the mean intensity values). To assess the contrast quantitatively in both sequences, we calculated the GM-WM ratios as follows: 1-cGM/scNAWM per lobe; 2-thalamus/posterior limb of the internal capsule; 3-Head of the caudate nucleus, putamen, globus pallidus (external and internal)/anterior limb of the internal capsule; 4-claustrum/WM of the insula; 5-amygdala, three regions of the hippocampus/WM of the temporal lobe; 6-three regions of the corpus callosum, two regions of the fornix/head of the caudate nucleus.

**Table 2.**
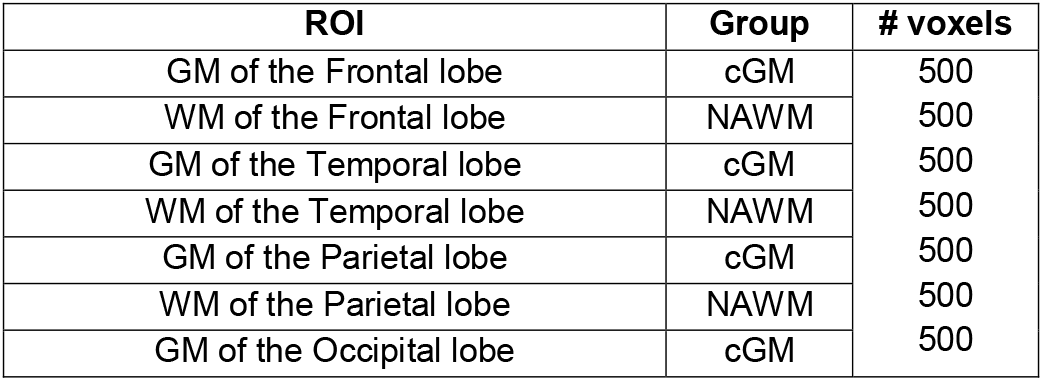
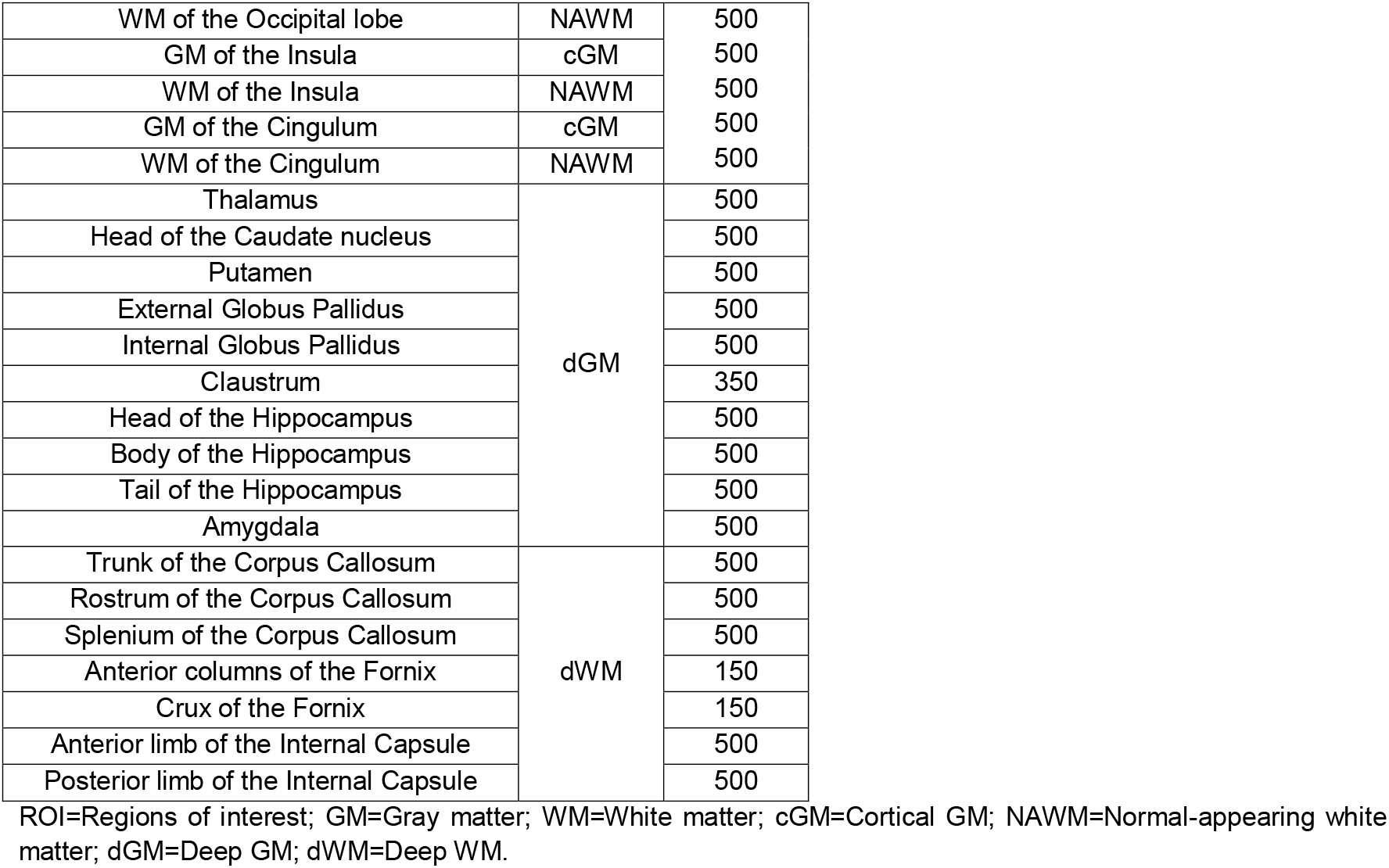
Regions of interest for quantitative analysis.

### Statistical analysis

Qualitative (i.e., categorical) variables were compared across the three fixative groups using Chi-square. Qualitative variables were also compared across the different FL, since T1 and T2 contrast is known to change longitudinally through fixation [20, 21] Quantitative variables were compared across the three fixatives using ANOVA or Kruskal-Wallis according to their distribution. All statistical tests were conducted using IBM SPSS statistics (version 29.0.2.0).

## Results

### Qualitative assessment

The inverted cGM-WM contrast was significantly different on T1w images across the different solutions (p<0.001). In AFS-fixed brains, all hemispheres showed non-inverted cGM-WM contrasts in most of the ROIs, while the other two solutions always showed inverted cGM-WM contrast in all of the ROIs assessed (p<0.001, Figure 2A). We found no significant difference in the frequency of inverted cGM-WM contrast in relation to the FL (Figure 2B). Out of the 6 AFS-fixed brains, all frontal, all cingulum, 4 insula, 3 parietal, 2 occipital and 3 temporal lobes showed non-inverted cGM-WM contrast, while 3 dGM-dWM showed non-inverted contrast, but this was not statistically significant (Figure 2C).

**Figure 2.**
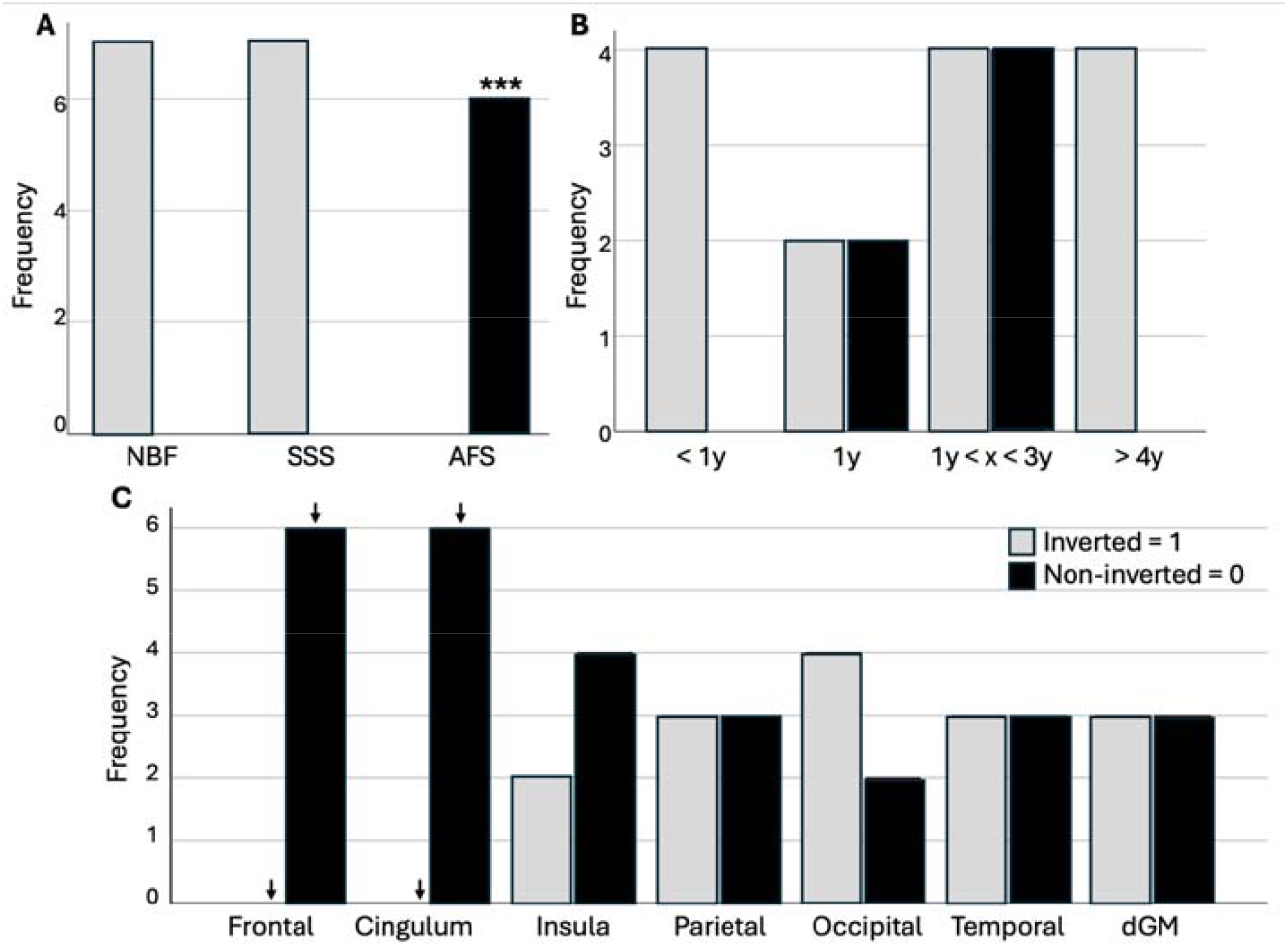
Qualitative assessment of the inverted contrast on T1w images. A) According to the fixative. B) According to the fixation length. C) In the different lobes and dGM of the AFS-fixed brains. NBF=Neutral=buffered formalin; SSS=Salt-saturated solution; AFS=Alcohol-formaldehyde solution; y=years. ↓ (before Bonferroni) 0.0055 < p < 0.05 ; *** p value < 0.001 after applying a Bonferroni correction.

When assessing this variable on T2w images, the contrast was never inverted, so there was no significant difference across the different solutions, nor across different FL.

As for the cGM-WM contrast sharpness on T1-weighted images, we found that the NBF-fixed brains showed a higher number of specimens with a sharp cGM-WM contrast (67.3% of all cases, p<0.0001), and fewer hemispheres with Low contrast (24.5% of all cases, p=0.01) or No contrast (8.2% of all cases, p<0.01) when compared with the other two solutions (Figure 3A). Furthermore, we found fewer cases with a sharp cGM-WM contrast in brains fixed with AFS (16.7% of all cases, p<0.001) (Figure 3A). We also found a significantly lower number of specimens with a sharp contrast in SSS-fixed brains (22.4% of all cases, p=0.01), and a significantly higher number of cases with No contrast in AFS specimens (35.7% of all cases, p<0.05), but these findings did not retain significance after a Bonferroni correction (Figure 3A).

**Figure 3.**
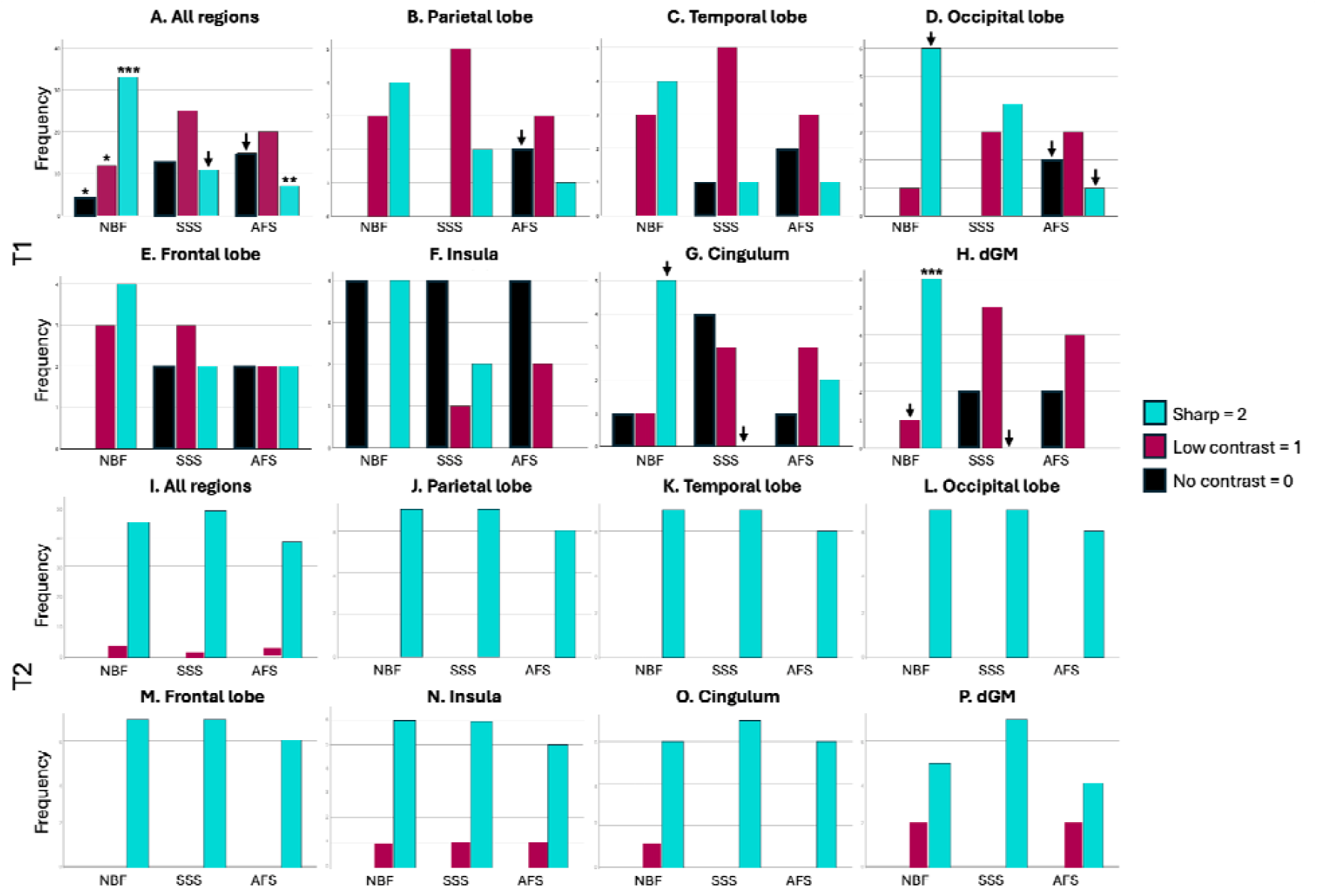
Qualitative assessment of the cGM-WM contrast sharpness in relation to the fixative solution. Top two rows show the cGM-WM contrast on T1w images, while bottom two rows represent the qualitative assessment on T2w images. dGM=Deep gray matter; NBF=Neutral-buffered formalin; SSS=Salt-saturated solution; AFS=Alcohol-formaldehyde solution. ↓ (before Bonferroni) 0.0055 < p < 0.05 ; * 0.001 < p < 0.0055 ; ** 0.0001 < p < 0.001 ; *** p < 0.0001.

The cGM-WM contrast sharpness was also assessed in different regions of interest (i.e., parietal, frontal, temporal, occipital, insular lobes, cingulum, and dGM nuclei). We found no significant differences in the temporal, frontal, and insular lobes. We found more hemispheres with No contrast in AFS-fixed brains in the parietal lobe, but this did not retain significance after a Bonferroni correction (p<0.05) (Figure 3B). In the occipital lobe, we found more cases with a sharp contrast in hemispheres fixed with NBF (p<0.05) and fewer in brains fixed with AFS (p<0.05), while more showed No contrast (p<0.05), but none of this retained significance after a Bonferroni correction (Figure 3D). In the cingulate gyrus, SSS-fixed hemispheres never showed sharp contrast (p<0.05), while most of the hemispheres fixed with NBF showed a sharp contrast (p=0.01), but this did not retain significance after a Bonferroni correction (Figure 3G). In the dGM, we found a significantly higher frequency of sharp contrast in the brains fixed with NBF (p<0.0001), while none was observed in SSS (p<0.05) nor AFS-fixed specimens (Figure 3H).

Finally, we found no significant differences of cGM-WM contrast sharpness on the T2w images. The cGM-WM contrast is mostly sharp in all the regions, while no cases of No contrast in any of the regions were observed (Figure 3I-P).

When we assessed the cGM-WM contrast sharpness in relation to the FL, we found overall no significant differences on T1w and T2w images. However, before a Bonferroni correction, there were differences in the insula (Figure 4F), cingulum (Figure 4G) and dGM nuclei (Figure4H) on T1w, as well as in all regions in the >4 years group (Figure 4I) and the cingulum in the >4 years group (Figure 4O) on T2w images. T1w and T2w images of the different specimens are shown in Figure 5.

**Figure 4.**
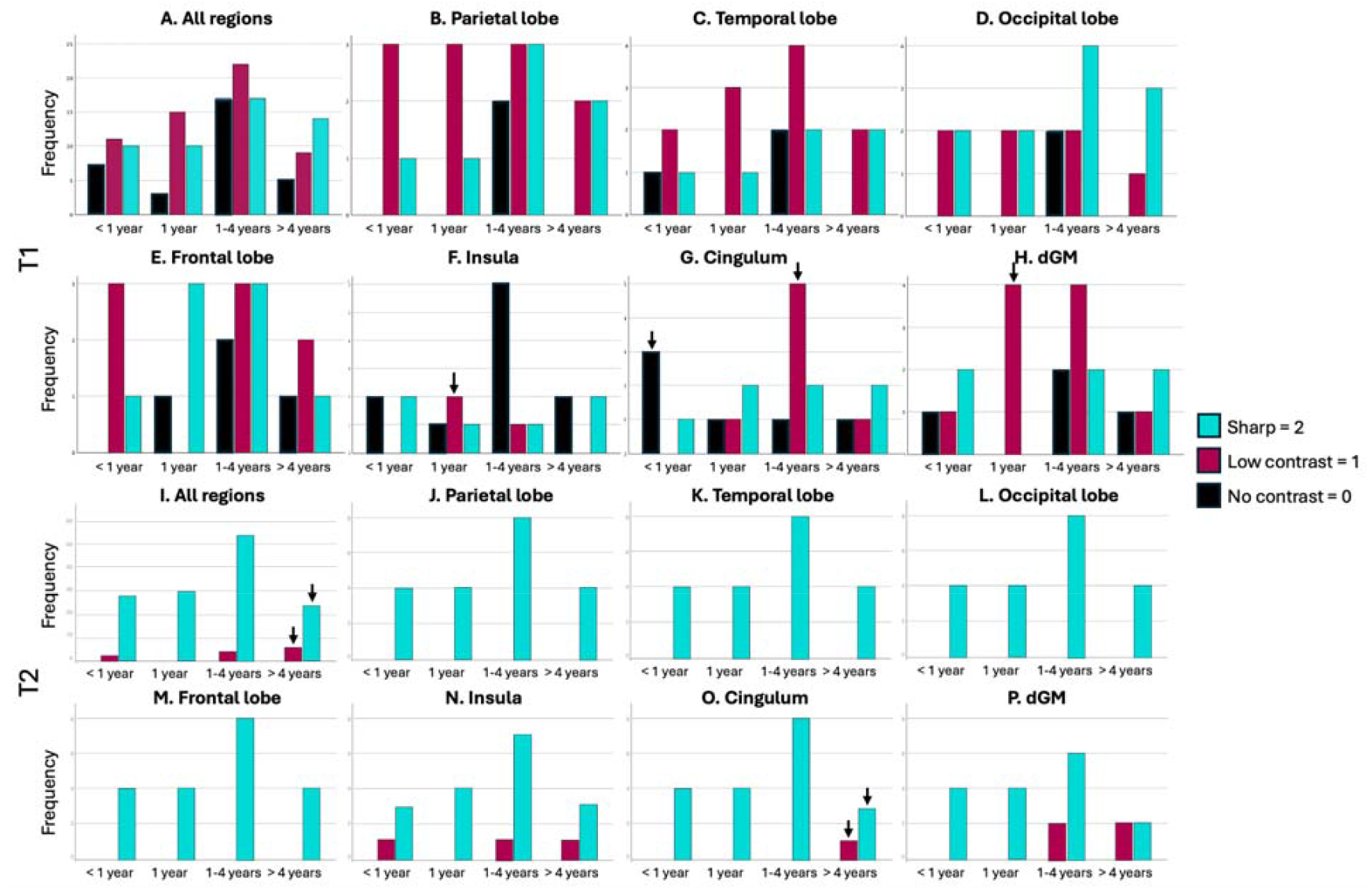
Qualitative assessment of the gray matter to white matter contrast sharpness in relation to the fixation delay. Top two rows show the GM-WM contrast in the T1-weighted sequence, while bottom two rows represent the qualitative assessment in T2-weighted images. dGM=Deep gray matter ↓ (before Bonferroni) 0.0055 < p < 0.05.

**Figure 5.**
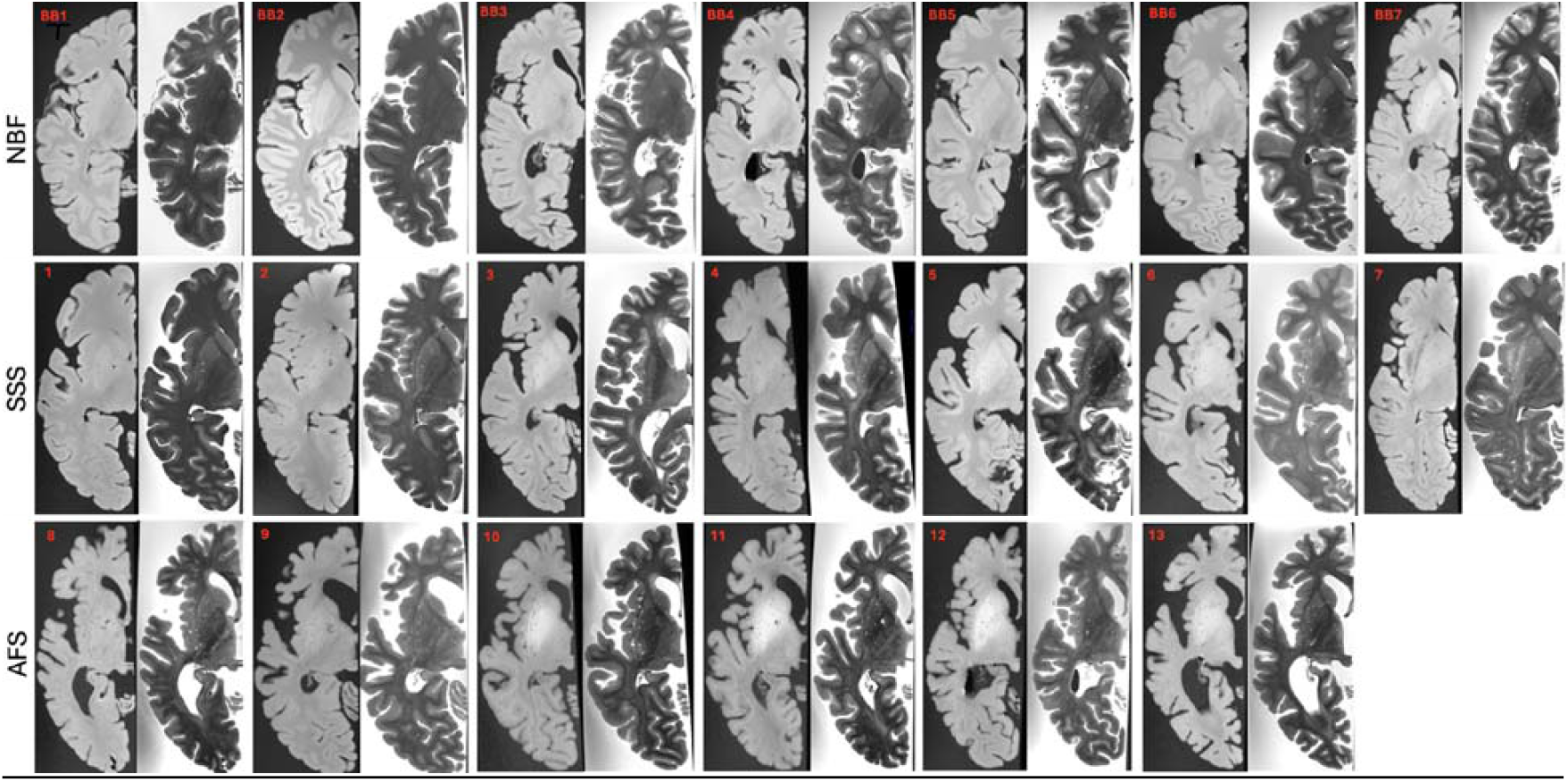
Qualitative assessment of the gray matter to white matter contrast in co-registered T1w and T2w images of all specimens. Top row shows the 7 NBF-fixed specimens, middle row shows the 7 SSS-fixed specimens and bottom row shows the 6 AFS-fixed brains. IDs are indicated in red.

### Quantitative assessment

Regarding the intensity ratios on T1w, we found that the AFS-fixed brains led to significantly lower ratios of GM-WM than those found in NBF-fixed brains in the frontal, temporal, parietal, occipital, insular and cingulate lobes. GM-WM ratios were also significantly lower in AFS-fixed brains than in SSS-fixed brains in the temporal, parietal, occipital, insular and cingulate lobes (Figure 6, top row, Table 3). Table 3 only shows the GM-WM intensity ratios that were found statistically significant, as shown in Figure 6. Also, Figure 1 of the Supplemental material shows per group mean intensities (cGM, NAWM, dGM, dWM) across the three fixatives.

**Table 3.**
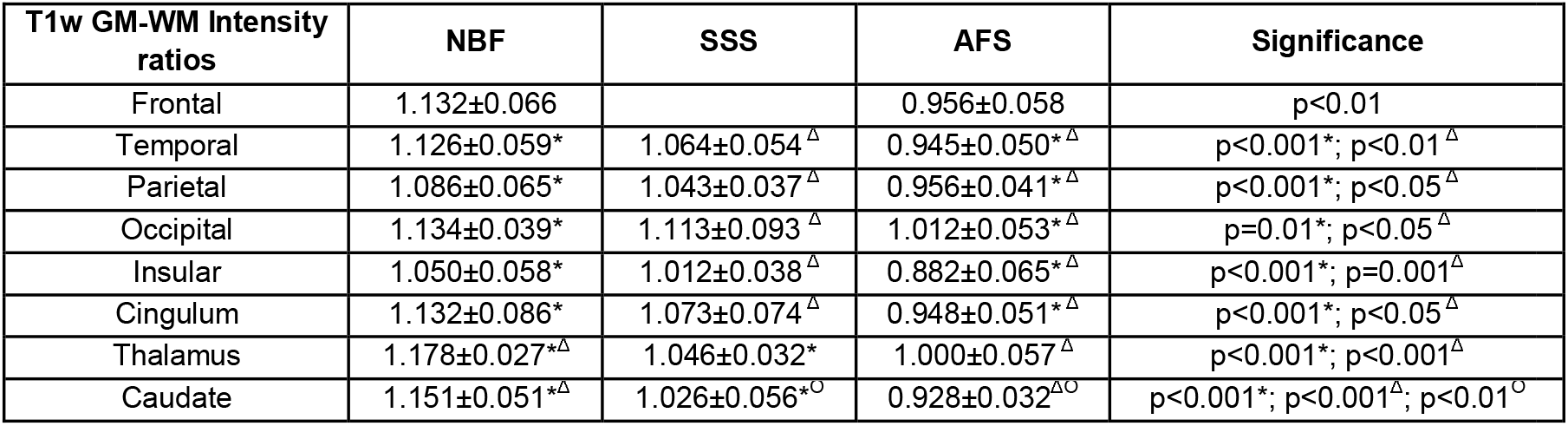
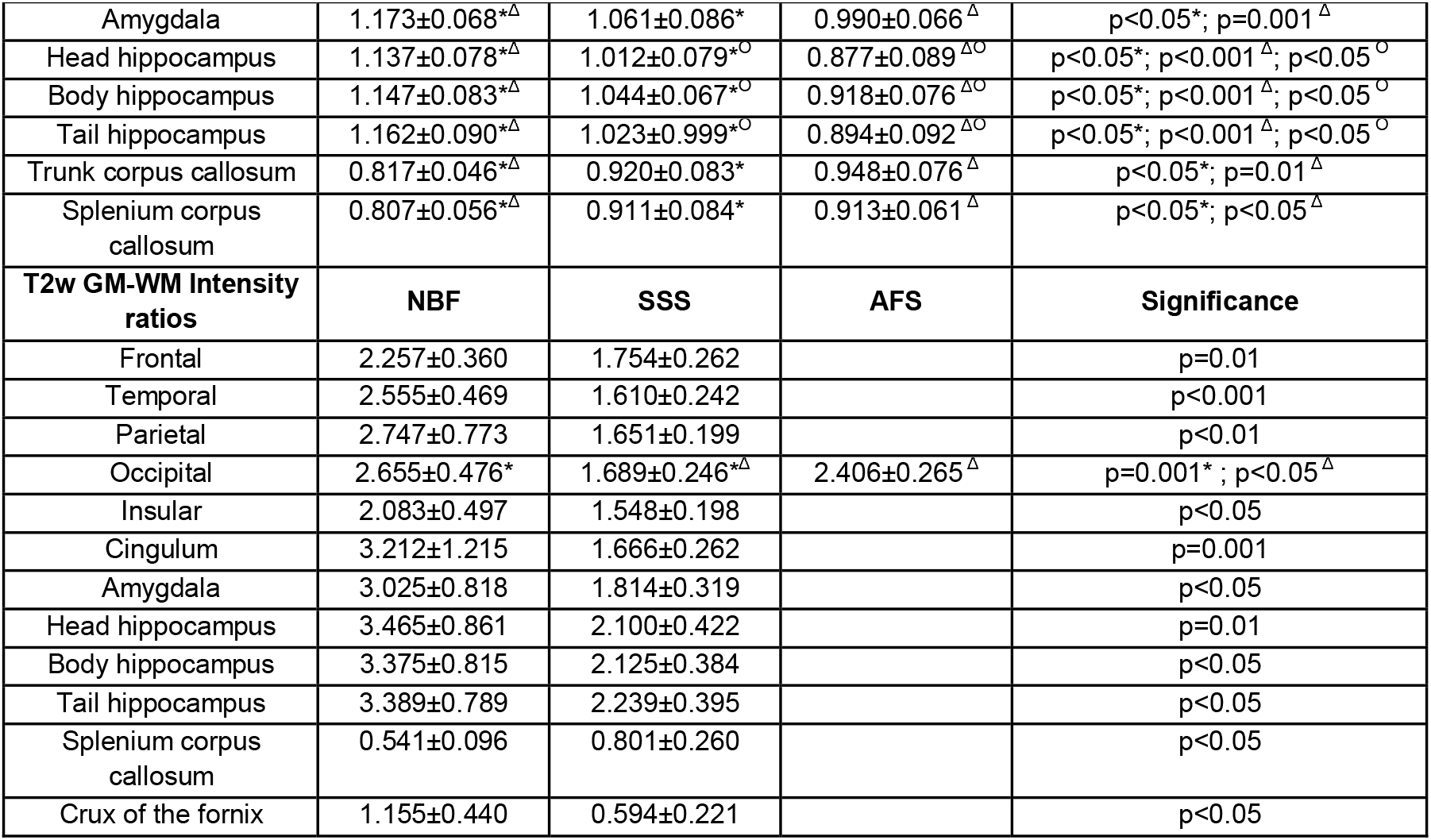
GM-WM mean intensity ratios of each region in T1wand T2w images.

**Figure 6.**
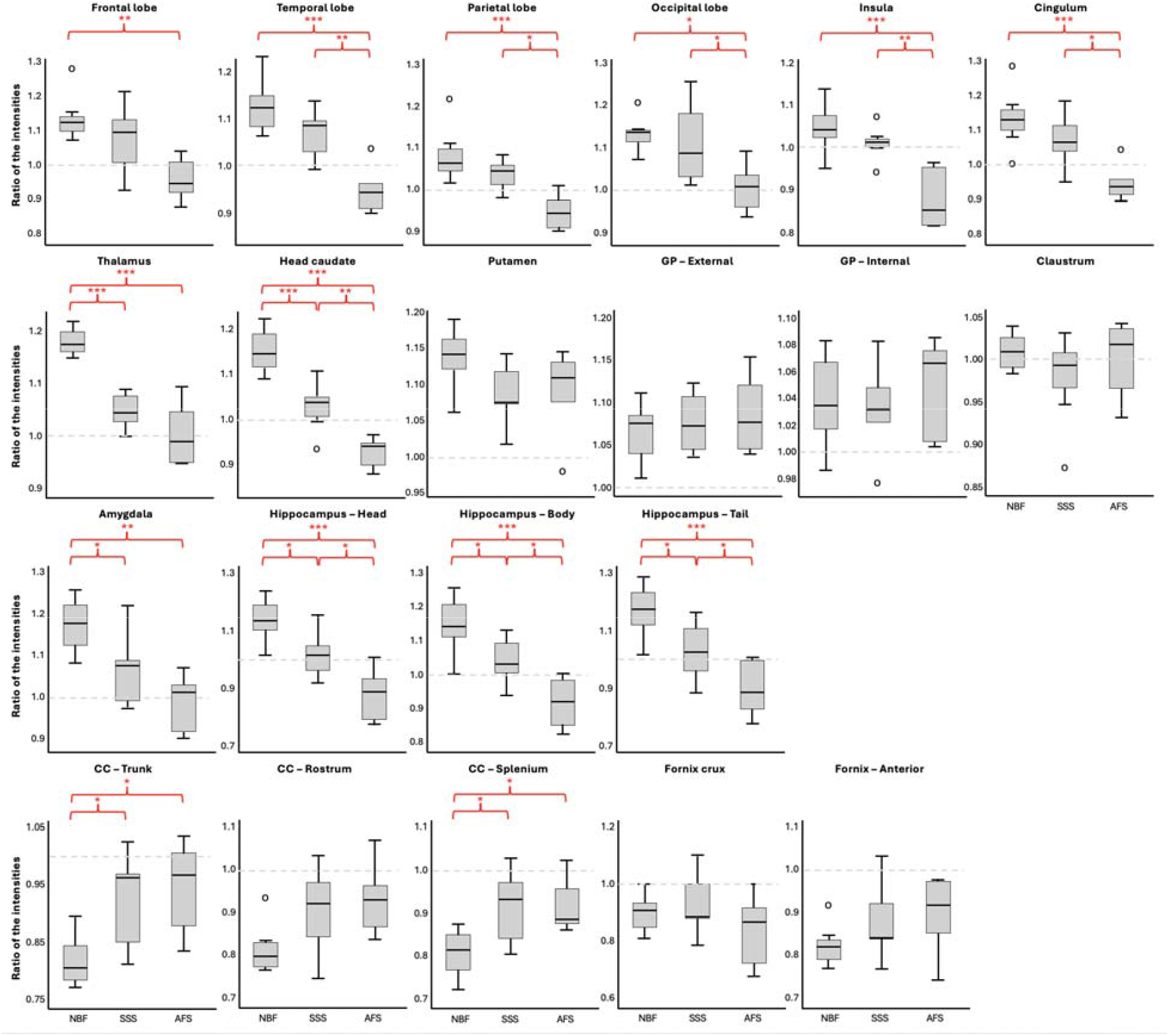
GM and WM T1w intensity ratios for each region of interest. The top line shows the ratios between GM and WM for each lobe. Second line shows the GM to WM ratio of the dGM nuclei (posterior limb of the internal capsule for thalamus, anterior limb of the internal capsule for head of the caudate, putamen and both globus pallidus, and white matter of the insula for claustrum). The third line shows the GM to WM ratios of the amygdala and hippocampus with the temporal WM (all four were considered dGM as well). Finally, the last line shows the WM to GM ratios of the dWM structures with the head of the caudate. GP=Globus pallidus; CC=Corpus callosum; NBF=Neutral-buffered formalin; SSS=Salt-saturated solution; AFS=Alcohol-formaldehyde solution. *0.01 < p < 0.05 ; **0.001 < p < 0.01; ***p < 0.001 after a Bonferroni correction.

In dGM regions, we found that the T1w intensity ratios of the thalamus of AFS-fixed brains was significantly lower than in NBF-fixed. Ratios of the SSS-fixed brains were also significantly lower than in NBF-fixed brains. We also found that the intensity ratios of the head of the caudate nucleus of AFS-fixed brains were significantly lower than in NBF-fixed brains and SSS-fixed brains. SSS-fixed brains also showed significantly lower caudate intensity ratios than NBF-fixed brains. Finally, we found no significant difference in the intensity ratios of the putamen, globus pallidus nor the claustrum in brains fixed with the three fixatives (Figure 6, second row).

We also found that the intensity ratios of the amygdala were significantly lower in brains fixed with AFS and SSS than in brains fixed with NBF. The intensity ratios of the three regions of the hippocampus were significantly lower in brains fixed with AFS and SSS than in brains fixed with NBF, and lower in AFS-fixed brains than in SSS (Figure 6, third row).

Finally, the intensity ratios of the trunk of the corpus callosum were significantly higher in brains fixed with AFS and SSS than in brains fixed with NBF. No significant difference was found for the rostrum of the corpus callosum, but we found similar differences of the intensity ratios of the splenium of the corpus callosum, where intensity ratios of AFS-fixed brains and SSS-fixed brains were significantly higher than in NBF-fixed brains (Figure 6, bottom row).

Regarding the T2w mean intensity ratios, we found that cGM-WM ratios of all lobes were significantly lower in brains fixed with SSS than in brains fixed with NBF. Furthermore, in the occipital lobe, SSS-fixed brains also showed significantly lower cGM-WM ratio than in brains fixed with AFS (Figure 7, top row).

**Figure 7.**
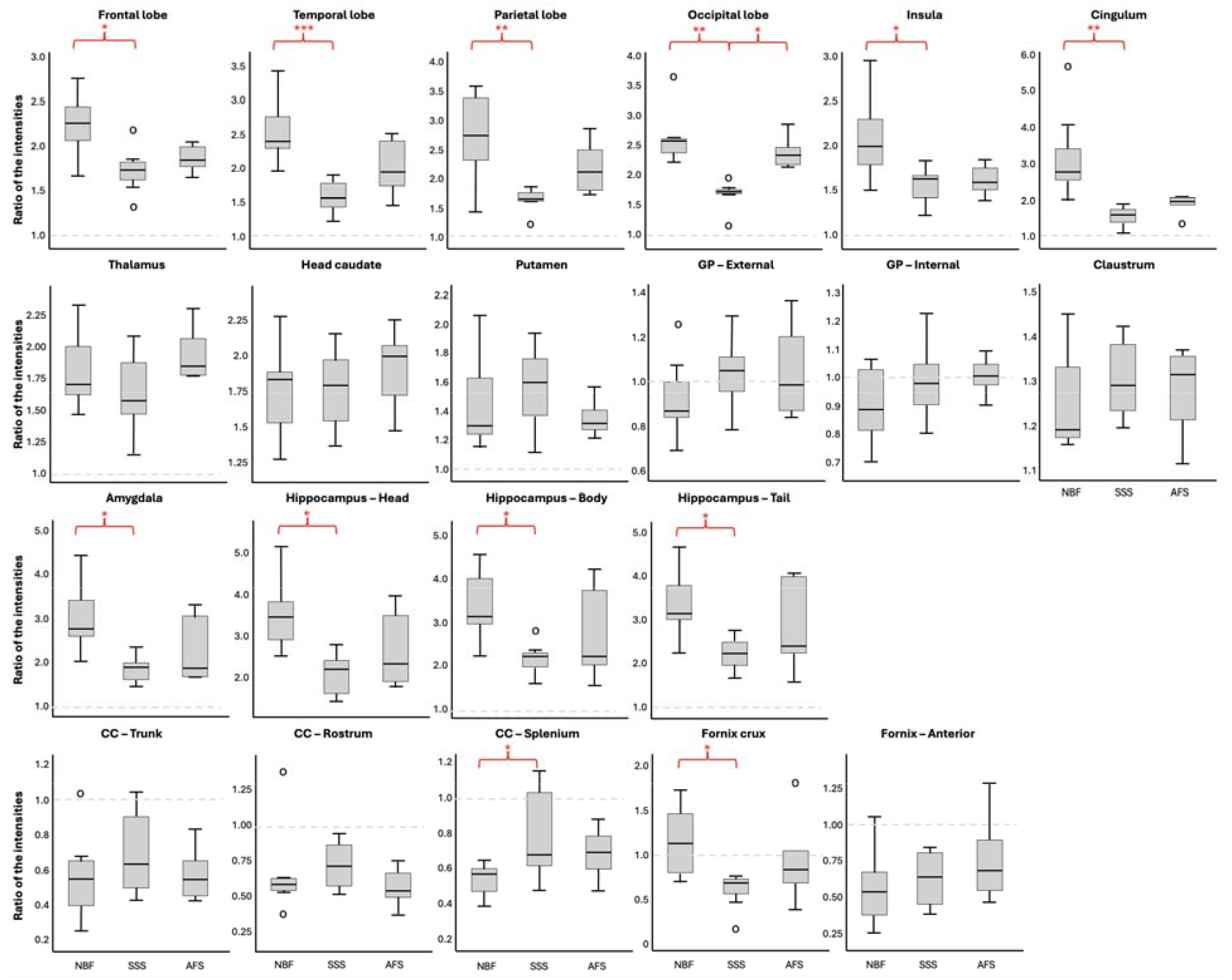
GM and WM T2w intensity ratios for each region of interest. Top line shows the ratios between GM and WM for each lobe. Second line shows the GM to WM ratio of the dGM nuclei (posterior limb of the internal capsule for thalamus, anterior limb of the internal capsule for head of the caudate, putamen and both globus pallidus, and white matter of the insula for claustrum). Third line shows the GM to WM ratios of the amygdala and hippocampus with the temporal WM (all four were considered dGM as well). Finally, last line shows the WM to GM ratios of the dWM structures with head of the caudate. GP=Globus pallidus; CC=Corpus callosum; NBF=Neutral-buffered formalin; SSS=Salt-saturated solution; AFS=Alcohol-formaldehyde solution. *0.01 < p < 0.05 ; **0.001 < p < 0.01, ***p < 0.001 after a Bonferroni correction.

The T2w dGM-WM mean intensity ratios (i.e., ratios of thalamus, head of the caudate nucleus, putamen, globus pallidus and claustrum) were not significantly different across the three solutions (Figure 7, second row). However, we found that the T2w mean intensity ratios of the amygdala and the three regions of the hippocampus were significantly lower in brains fixed with SSS than in brains fixed with NBF (Figure 7, third row).

Finally, as for the dWM ratios, we found no significant difference in the trunk and the rostrum of the corpus callosum, nor in the anterior column of the fornix. However, we found that the ratios of the splenium of the corpus callosum of the SSS-fixed brains were significantly higher than in NBF-fixed brains. Furthermore, we found that the ratios of the crux of the fornix were significantly lower in brains fixed SSS than with NBF.

## Discussion

In this study, we systematically assessed the impact that fixation by three different fixative solutions exerts on T1w and T2w MRI tissue contrasts in different structures of the human brain. First, we examined whether GM-WM contrast was inverted in T1w images, leading to the GM appearing as hyperintense in relation to the WM (i.e. as opposed to the contrast profiles of *in vivo* T1w images showing hypointense GM in relation to WM). Previous studies suggest that post-mortem brains fixed with NBF show different T1w contrasts because of fixative distribution into the tissue [10, 29]. Additionally, T1w GM-WM contrast and T1 relaxation times were measured in five formalin-fixed human brains as a function of time [23]. They showed that a reversal of the contrast was observed around the 4^th^ day of immersion, which was in relation to a greater decrease of the GM relaxation times than in WM. Based on this work, our initial hypothesis was that the T1w images of our anatomy laboratory hemispheres might not always show an inverted GM-WM contrast, since both anatomic solutions have a much lower proportion of formaldehyde (0.8% in the SSS and 1.4% in the AFS compared to 4% in NBF). We also considered that the FL could impact the reversal of the signal, and that this would be more evident in specimens with a longer FL.

In T1w images, only AFS-fixed brains showed non-inverted contrast in some lobes (all cases). All SSS and NBF brains showed an inverted contrast, comparable to what other studies have shown in NBF cases [10, 29]. We suspect that this lack of inversion in some regions of our AFS specimens may be related to two phenomena: 1-changes in AFS may be slower than changes produced by SSS or NBF, and/or 2-distribution and diffusion of the fixative might not have been sufficiently homogeneous in the AFS-fixed specimens of our sample. Regarding the first consideration, based on our prior examinations, AFS-fixed bodies require between 8 months to 1 year to be deemed macroscopically well fixed for dissection and teaching purposes (according to the texture, color, and humidity of the body). In contrast, SSS-fixed bodies only take 6 to 8 weeks to be usable in dissections, while NBF brains are immersed for only 3 weeks prior to use [10]. Consequently, we examined the inversion of the contrast in relation to the FL, but did not find statistically significant associations. However, only 2 AFS-fixed specimens had a FL of 3 years, and those showed more areas with inverted contrast (cases 8 and 9 in Figure 5 and Table 1). It is therefore possible that a larger sample of specimens with longer FLs would show a relationship between FL and inversion of the GM-WM contrast in the AFS-fixed specimens. Regarding inhomogeneity in distribution of the fixative, our previous study on T1w and T2w contrasts of suboptimally fixed *in situ* human brains [34] found a non-inverted contrast on T1w images. When extracting the brains and sectioning the hemispheres, we found remaining blood in the leptomeninges, the cortex, and the deeper WM, where the non-inverted contrasts on the T1w images were observed. In some cases, these areas corresponded to the vascular territories of arteries that were visibly obstructed by clots, so it is conceivable that the fixative could not be fully and homogeneously distributed in all the tissue [34]. In the present sample of AFS-fixed brains, we have not observed any macroscopical accumulation of blood affecting the cortex or dWM, so we did not systematically inspect the vessels of these specimens. Still, it is always possible that smaller vessels are affected by obstructions (i.e. small vessels disease, especially in our older population) and this could be related to inhomogeneous fixation and reversal of the signal only in some lobes.

Regarding the anatomical location of non-inverted contrast in the AFS-fixed brains, we found that all frontal and cingulum lobes were non-inverted, while 4 insula, 3 parietal, 3 temporal, and only 2 occipital lobes out of 6 specimens showed a non-inverted contrast. We hypothesize that the difference in the inversion of the contrast across the ROIs could be related to vascular and myelin anterior-to-posterior differences in the brains. First, the brains are perfused through whole body perfusion by the injection of the fixative into the common carotid artery (at its proximal part into the neck). The vertebral system, responsible for the fixation of the occipital lobe and posterior regions of the brain through the posterior cerebral arteries, is less affected by clots than the carotid system, responsible for the perfusion of the rest of the brain through the middle and anterior cerebral arteries [35, 36]. Because the injection of the fixative solution, using a pump, comes from the carotid artery and has a pressure of 310 mmHg, this could push the clots towards the distal branches of the carotid system instead of the vertebral system, potentially determining a more heterogeneous distribution of the fixative in the anterior compared to the posterior territories of the brain. Second, there is a fronto-occipital (antero-posterior) gradient of myelination in the development of the brain [37], leading to fundamental differences in the myelin composition in the different ROIs of the hemispheres, especially in the frontal lobe that loses myelin with aging [37]. Indeed, a previous study hypothesized that the myelin phospholipid structure was decomposed by formalin fixation, which somewhat counteracts the general reductive effect of the fixation procedure on relaxation times [23]. Another study suggested that myelin is lost in the frontal lobe and superior regions of the brains with aging [37]. Therefore, our results are in line with the fact that the inversion of the T1w contrast could be related to differential anterior-to-posterior changes in the myelin. Finally, in a previous report of our team that scanned one NBF-fixed in situ brain once a month (starting two months after the injection of the fixative [38], also see Figure 3 in the Supplemental materials), we found that the inversion of the cGM-WM contrast started in the occipital lobes (starting from the second visit (3 months) after injection of the fixative), followed by the parietal lobe (starting from the third visit (4 months)) and finally by the frontal lobe (from the 5^th^ visit) (i.e., from posterior to anterior), reflecting the same differences in the inversion of the GM-WM contrast as we see in the AFS-fixed specimens of this study.

In T2w images, we found that the contrast was never inverted in the brains fixed with the three fixatives, reflecting that T2w characteristics are not affected by the chemical fixation as it is the case on T1w images. Therefore, post-mortem T2w images resemble those *in vivo* [10, 30], translating the suitability of brains fixed in anatomy laboratories for using T2w MRI sequences.

We assessed this variable because differences in the sharpness of the GM-WM contrast could influence studies that use segmentation of different anatomical structures. Indeed, manual segmentation by experts or automatic segmentation by softwares are highly dependent on the intensity of voxels at the boundary between two different tissue types and the partial volume of each tissue in boundaries’ voxels [39-41]. This is important for example in MRI studies interested in the volume of different regions and nuclei, where differences in the fixative used could impact the extracted volumes for the same structures [39, 40]. This is also important since automated segmentation methods are inevitably impacted by tissue contrast and would have been biased if we hadn’t used manual segmentation. Therefore, we have used neuroanatomy expertise to accurately segment small regions (at comparable levels in each scan) to show how their respective intensities were impacted by the fixation process and the different fixatives.

In T1w images, out of 7 ROIs per specimen (frontal, temporal, parietal, occipital, cingulum, insular lobes and dGM), we found that NBF-fixed brains had the lowest proportion of ROIs without contrast: only 8.2% compared to 26.5% in SSS- and 35.7% in AFS-fixed brains. Also, SSS- and AFS-fixed specimens showed the highest proportion of ROIs with a Low contrast: 51% and 47.6%, as opposed to 24.5% of the NBF-fixed brains’ ROIs. Finally, NBF-fixed cases also showed the highest proportion of sharp contrast: 67.3%, while only 22.4% of SSS- and 16.7% of AFS-fixed brains’ regions showed Sharp contrast. This reflects the lowest GM-WM contrast in AFS-fixed brains, followed by SSS-fixed specimens and that the highest quality in terms of sharp boundaries is provided by NBF-fixed brains.

The occipital lobe and dGM were the ROIs that showed the highest differences in the proportion of cases with sharp boundaries across the three solutions. Occipital lobes showed sharper contrasts in NBF-fixed brains, which might be related to the occipital cortex being more compact and with less gyrification [42-44]. In fact, a faster fixation by immersion in the cortical surface by NBF may be responsible for a sharper contrast. Plus, even in the AFS specimens, we found that the occipital lobes were the least affected ROI by the non-inverted contrast, which points to the intrinsic characteristics of the area of the brain to explain a faster, more homogeneous fixation regardless of the solution employed. Interestingly, in the dGM, we never found sharp contrast (which happened to be highly significant), except in NBF fixed brains, even though the fixative had to diffuse from the surface, as opposed to the specimens fixed with SSS and AFS, where the fixative gets to the tissue through the vessels. We think that this may be related to the anatomical solutions having a much lower concentration of formaldehyde (0.8% for SSS and 1.4% for AFS) than the NBF solution (4% formaldehyde).

In T2w images we found no difference in the sharpness of the GM-WM borders between the three fixatives, where almost all ROIs in all specimens showed a sharp contrast. This is in accordance with previous studies that showed that *ex vivo* T2w quality looked similar than *in vivo* T2w images, highlighting this sequence suitability for *ex vivo* research using MRI [10, 22, 30, 34].

Finally, we also considered the possibility that a lower contrast sharpness could reflect an incomplete fixation process that could eventually be completed with a longer exposure to the fixative. Therefore, we assessed the GM-WM contrast sharpness in relation to the FL, but we found no significant difference in any of the ROIs, neither in T1w or T2w images. However, since our sample size is small and we have no AFS-fixed brains with a FL longer than 3 years, we cannot unequivocally conclude on the effect of FL on the AFS-fixed brains T1w contrast, which seems to be the most heterogeneous or incomplete of the three solutions.

### Voxel-wise intensity ratios

It has been shown that MRI properties change after fixation in NBF [10-12, 20-24, 26, 29, 45-47], so we wanted to assess if the intensity of the voxels fixed with the other two solutions would also be affected. Furthermore, we wanted to corroborate what was observed in our qualitative assessment, so we extracted intensities in 29 ROIs of cGM, scNAWM, dGM and dWM to calculate ratios that would reflect the GM-WM contrast. Indeed, a ratio closer to 1 reflects a lower contrast (i.e., since it shows that GM and WM intensity values are similar) while a ratio further from 1 reflects a better GM-WM contrast. Additionally, a ratio lower than 1 would reflect a non-inverted contrast (i.e., WM hyperintense with a higher numeric value in the denominator than GM in the numerator).

In T1w images, we found that AFS-fixed brains had lower ratios than NBF- and SSS-fixed brains in all the lobes. They also had lower ratios in the thalamus, caudate, amygdala and hippocampus. Also, the ratios of the trunk and the splenium of the corpus callosum were higher in AFS- and SSS-fixed brains than in NBF-fixed brains. These differences in the ratios of AFS-fixed brains are related to a higher intensity of WM (either NAWM or dWM) (Supplemental Table 1 and Figure 1), which is related to the lack of inversion of the contrast. Indeed, in all the lobes and dGM, the mean ratios of AFS-fixed brains are lower than 1, and when calculating the difference between the GM-WM ratios and 1 (1 representing no-contrast), we observed negative values, translating that the contrast was not completely reversed. We also obtained values closest to 0, translating the lowest contrast (Ratios of 1 minus 1) of the three solutions (Supplemental Figure 2). A potential explanation for these findings in AFS-fixed brains is that the multiple chemicals included in the AFS (i.e., high concentrations of ethanol, glycerol and phenol, as well as Dettol, a detergent used as a degreaser) have a distinct effect on the WM. We hypothesize that the AFS solution affects the myelin of the WM in its own idiosyncratic way determining a more heterogeneous contrast on MRI compared to the NBF and SSS solutions. Indeed, our hypothesis is supported by the fact that AFS-fixed brains have lower quality of Luxol fast blue and Bielschowsky stains (i.e., techniques that stain myelin and axons) [18], that myelin water fraction and relaxation times are generally modified with fixation [48], and that phospholipids of the myelin sheet may be degraded by the fixation process, but could be modified in delipidization processes [49].

In T2w images, we found that SSS-fixed brains showed lower GM-WM intensity ratios in all the lobes, in the amygdala and in the hippocampus. Again, Supplemental Table 2 and Figure 1 show that the WM intensities change more than GM regions. However, in SSS-fixed brains, GM regions were also higher, leading into overall intensity ratios that were lower. As observed in the qualitative assessment, this did not affect the sharpness of the GM-WM contrast, so we suggest that this is probably due to the overall higher intensities in these brains, since sodium (i.e., present in high concentration in the SSS-fixed brains) is known to produce hyperintense signal on MRI [50]. In Supplemental Figure 2, we show that the difference between the mean ratios of SSS-fixed brains and the value 1 are higher than 0, reflecting a sharp GM-WM contrast, as also observed in the qualitative assessment.

### Limitations

We acknowledge that our study isn’t without limitations. We used a convenience sample of specimens that we tried to match in relation to confounding variables (i.e., age at the time of death), but we still found ourselves with older specimens in the groups fixed in the anatomy laboratory (i.e., SSS- and AFS-fixed brains). This is a consequence of the overall older ages of the donors of our anatomy laboratory, which is beyond our control. We were also unable to include AFS-fixed brains fixed for more than 4 years, but we are working on building a larger sample with longer FL for future studies. Finally, our small sample size limits the statistical analysis approaches that we could use, which is why we qualitatively assessed the structural MRIs and avoided attempting regressions with multiple covariables that demand much larger samples.

### Conclusion

We conclude that T1w images are more affected by the fixation process, inverting the contrast of *in vivo* T1w and reducing the GM-WM contrast in general, while T2w images remain similar to *in vivo* scans and maintain a sharp contrast. AFS-fixed brains seem to be more affected by longitudinal relaxation (T1), while SSS-fixed brains seem more affected by the transversal relaxation (T2). Furthermore, the differences in the intensity ratios across fixatives seem to be related to more marked changes in the WM intensity values across fixatives, not GM values, reflecting the importance of the changes of the myelin during the fixation process. Overall, brains fixed in anatomy laboratories could be used for MRI studies and the T2w sequence seems to be the most adequate for structural analyses in different brain regions.

## Acknowledgements

We would like to thank the generosity of the body donors and their families for making our projects possible. We also acknowledge funding resources and the anatomy laboratory staff of the Université du Québec à Trois-Rivières for their support and help in the lab. We would also like to thank the Cerebral Imaging Center staff for their help in managing the scanning schedule. All the authors are funded by the Government of Canada | Natural Sciences and Engineering Research Council of Canada. MRI data collection was also funded by Dr. Dadar’s grant from the Canadian Institutes of Health Research.

## List of figures and tables

Supplemental Table 1. Mean intensities of each of the ROIs of T1w images.

Supplemental Table 2. Mean intensities of each of the ROIs of T2w images.

**Supplemental figure 1. Quantitative assessment of the mean intensity in each ROI group in relation to the fixative**. Left four graphs show the intensities in the T1-weighted sequence, while the four right graphs represent the assessment in T2-weighted images.

cGM=Cortical gray matter; NAWM=Normal-appearing white matter; dGM=Deep gray matter; dWM=Deep white matter; NBF=Neutral-buffered formalin; SSS=Salt-saturated solution; AFS=Alcohol-formaldehyde solution.

*0.01 < p < 0.05 ; **0.001 < p < 0.01; ***p < 0.001 after a Bonferroni correction.

**Supplemental figure 2. Quantitative assessment of the difference of the ratios with 1 (i.e**., **GM-WM ratio value minus 1) to reflect the GM-WM contrast**. A value of 0 reflects no contrast; the higher the value, the higher the contrast. A negative value reflects a non-inverted contrast. In T1w, we found that the values are closer to 0 than in T2w images, reflecting less contrast. In T1w, AFS-fixed brains show the lowest and non-reversed contrast, with negative values and closest to 0. In T2w images, SSS-fixed brains showed closest values to 0, reflecting less contrast than in the brains fixed with the other two solutions, but still further from 1 than in T1w, translating the overall higher contrast of T2w images.

NBF=Neutral-buffered formalin; SSS=Salt-saturated solution; AFS=Alcohol-formaldehyde solution.

*0.01 < p < 0.05 ; ***p < 0.001 after a Bonferroni correction.

**Supplemental figure 3. Longitudinal assessment of the GM-WM contrast of an *in situ* brain fixed by perfusion with NBF**. This figure was presented at the annual conference of the Society for Neuroscience in 2021 as a virtual poster presentation. First scan (visit 1) was done 2 months after the injection of NBF, which showed no inversion of the contrast (most of the tissue showed preserved GM-WM contrast, reflecting no inversion of the GM-WM). At visit 2, occipital lobes and periventricular spaces showed loss of GM-WM contrast, indicating the beginning of the inversion of the contrast. Inversion of the contrast was completed at visit 6 (8 months after the injection of NBF). (Frigon et al., 2021, Society for Neuroscience).

