## Supplementary material for "Comparison of T1- and T2-weighted MRI contrasts of *ex vivo ex situ* brains fixed with solutions used in gross anatomy laboratories"

Supplemental Table 1. Mean intensities of each of the ROIs of T1w images.

| **ROI** | **ROI group** | **Mean intensity NBF** | **Mean intensity SSS** | **Mean intensity AFS** | **Significance** |
| --- | --- | --- | --- | --- | --- |
| 1 | cGM | 90.238 | 90.176 | 90.538 | p>0.05 |
| 2 | NAWM | 79.851* | 84.551 | 94.903* | p=0.001* |
| 3 | cGM | 87.524 | 90.257 | 87.960 | p>0.05 |
| 4 | NAWM | 77.856*^Δ^ | 84.952*^O^ | 93.310^ΔO^ | p=0.024* ; p<0.001^Δ^ ; p=0.01^O^ |
| 5 | cGM | 87.651 | 87.955 | 88.200 | p>0.05 |
| 6 | NAWM | 80.868* | 84.420^Δ^ | 92.423*^Δ^ | p<0.001* ; p=0.008^Δ^ |
| 7 | cGM | 89.628 | 92.839 | 92.358 | p>0.05 |
| 8 | NAWM | 79.107* | 83.894 | 91.447* | p=0.004* |
| 9 | cGM | 94.383 | 95.814 | 90.306 | p>0.05 |
| 10 | NAWM | 90.029* | 94.748^Δ^ | 102.719*^Δ^ | p<0.001* ; p=0.008^Δ^ |
| 11 | cGM | 93.330 | 93.814 | 88.762 | p>0.05 |
| 12 | NAWM | 82.684* | 87.500 | 93.614* | p=0.002* |
| 13 | dGM | 91.063 | 88.980 | 94.632 | p>0.05 |
| 14 | dGM | 95.933 | 93.613 | 96.396 | p>0.05 |
| 15 | dGM | 94.820 | 99.189 | 114.024 | p>0.05 |
| 16 | dGM | 88.635* | 98.031^Δ^ | 113.167*^Δ^ | p<0.001* ; p=0.018^Δ^ |
| 17 | dGM | 86.594* | 94.235^Δ^ | 109.555*^Δ^ | p<0.001* ; p=0.013^Δ^ |
| 18 | dWM | 78.258*^Δ^ | 85.599*^O^ | 90.990^ΔO^ | p=0.001* ; p<0.001^Δ^ ; p=0.022^O^ |
| 19 | dWM | 78.241* | 83.991 | 89.410* | p<0.001* |
| 20 | dWM | 77.239*^Δ^ | 84.736* | 87.744^Δ^ | p=0.002* ; p<0.001^Δ^ |
| 21 | dWM | 77.350*^Δ^ | 85.213*^O^ | 94.707^ΔO^ | p=0.028* ; p<0.001^Δ^ ; p=0.01^O^ |
| 22 | dWM | 83.406* | 91.306^Δ^ | 104.085*^Δ^ | p<0.001* ; p=0.01^Δ^ |
| 23 | dGM | 91.125 | 89.811 | 92.204 | p>0.05 |
| 24 | dWM | 87.491* | 91.252 | 95.594* | p=0.006* |
| 25 | dWM | 86.067 | 87.094 | 80.846 | p>0.05 |
| 26 | dGM | 88.304* | 85.734 | 81.501* | p=0.026* |
| 27 | dGM | 89.100 | 88.453 | 85.422 | p>0.05 |
| 28 | dGM | 90.305 | 86.738 | 83.141 | p>0.05 |
| 29 | dGM | 91.011* | 92.863^Δ^ | 103.327*^Δ^ | p=0.013* ; p=0.038^Δ^ |

Supplemental Table 2. Mean intensities of each of the ROIs of T2w images.

| **ROI** | **ROI group** | **Mean intensity NBF** | **Mean intensity SSS** | **Mean intensity AFS** | **Significance** |
| --- | --- | --- | --- | --- | --- |
| 1 | cGM | 28.692 | 34.679 | 33.390 | p>0.05 |
| 2 | NAWM | 12.764* | 20.394* | 17.894 | p=0.005* |
| 3 | cGM | 24.411 | 29.693 | 27.750 | p>0.05 |
| 4 | NAWM | 9.732* | 18.642* | 14.418 | p=0.002* |
| 5 | cGM | 29.151 | 33.492 | 28.034 | p>0.05 |
| 6 | NAWM | 11.114* | 20.725*^Δ^ | 13.257^Δ^ | p<0.001* ; p=0.01^Δ^ |
| 7 | cGM | 23.207 | 26.322 | 24.286 | p>0.05 |
| 8 | NAWM | 8.948* | 15.515*^Δ^ | 10.237^Δ^ | p=0.003* ; p=0.021^Δ^ |
| 9 | cGM | 31.019 | 38.113 | 33.314 | p>0.05 |
| 10 | NAWM | 14.703*^Δ^ | 25.233* | 20.908^Δ^ | p<0.001* ; p=0.046^Δ^ |
| 11 | cGM | 36.639 | 39.165 | 37.316 | p>0.05 |
| 12 | NAWM | 12.033* | 24.463* | 19.257 | p<0.001* |
| 13 | dGM | 17.711*^Δ^ | 27.673* | 25.537^Δ^ | p=0.009* ; p=0.027^Δ^ |
| 14 | dGM | 20.113*^Δ^ | 32.912* | 28.780^Δ^ | p=0.002* ; p=0.046^Δ^ |
| 15 | dGM | 16.968* | 28.284*^Δ^ | 20.542^Δ^ | p=0.002* ; p=0.026^Δ^ |
| 16 | dGM | 10.655* | 19.571* | 15.718 | p=0.005* |
| 17 | dGM | 10.220 | 18.595 | 15.267 | p>0.05 |
| 18 | dWM | 10.852* | 22.459* | 16.581 | p=0.002* |
| 19 | dWM | 12.656* | 23.465*^Δ^ | 16.223^Δ^ | p<0.001* ; p=0.028^Δ^ |
| 20 | dWM | 10.797*^Δ^ | 25.383* | 19.537^Δ^ | p<0.001* ; p=0.01^Δ^ |
| 21 | dWM | 10.002* | 17.275* | 13.229 | p=0.002* |
| 22 | dWM | 11.587* | 19.183* | 15.457 | p=0.021* |
| 23 | dGM | 28.391 | 32.728 | 30.048 | p>0.05 |
| 24 | dWM | 10.950 | 20.359 | 21.670 | p>0.05 |
| 25 | dWM | 22.590 | 19.289 | 25.963 | p>0.05 |
| 26 | dGM | 32.197 | 37.644 | 35.221 | p>0.05 |
| 27 | dGM | 31.737 | 38.093 | 34.308 | p>0.05 |
| 28 | dGM | 32.104 | 40.326 | 36.972 | p>0.05 |
| 29 | dGM | 18.561* | 33.002* | 26.729 | p<0.001* |

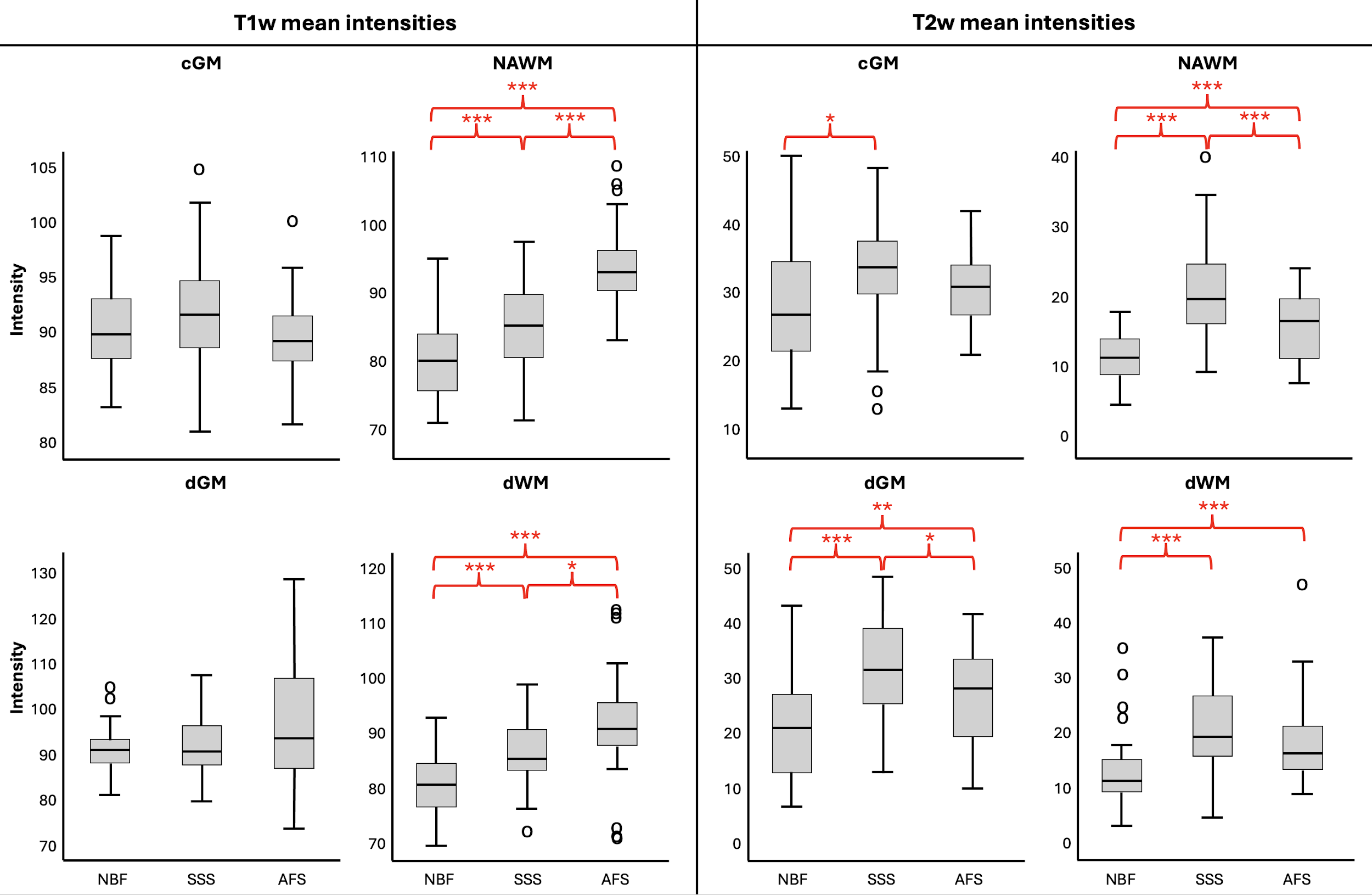

**Supplemental figure 1. Quantitative assessment of the mean intensity in each ROI group in relation to the fixative.** Left four graphs show the intensities in the T1-weighted sequence, while the four right graphs represent the assessment in T2-weighted images.

cGM=Cortical gray matter; NAWM=Normal-appearing white matter; dGM=Deep gray matter; dWM=Deep white matter; NBF=Neutral-buffered formalin; SSS=Salt-saturated solution; AFS=Alcohol-formaldehyde solution.

*0.01 < p < 0.05 ; **0.001 < p < 0.01; ***p < 0.001 after a Bonferroni correction.

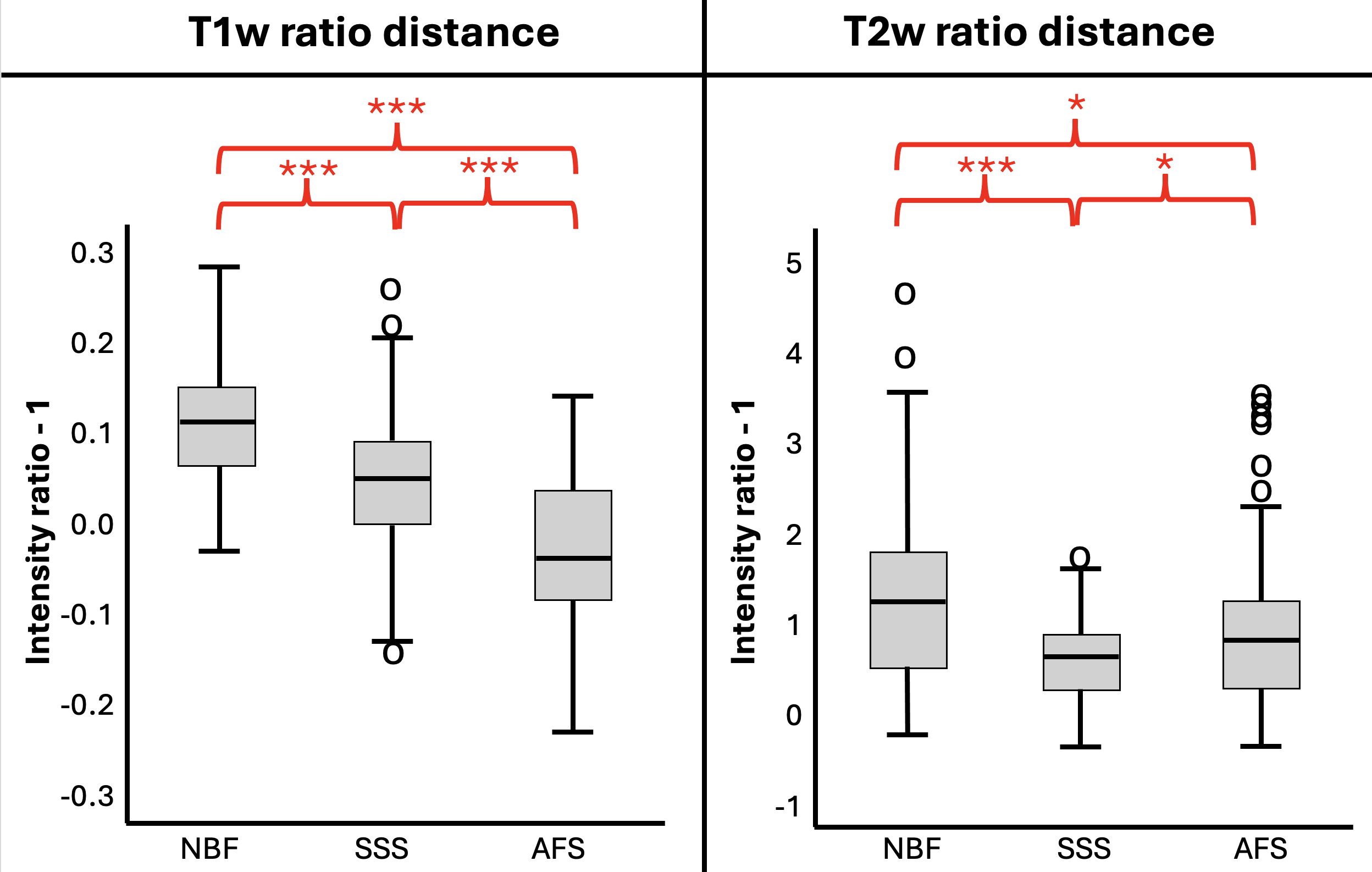

**Supplemental figure 2. Quantitative assessment of the difference of the ratios with 1 (i.e., GM-WM ratio value minus 1) to reflect the GM-WM contrast.** A value of 0 reflects no contrast; the higher the value, the higher the contrast. A negative value reflects a non-inverted contrast. In T1w, we found that the values are closer to 0 than in T2w images, reflecting less contrast. In T1w, AFS-fixed brains show the lowest and non-reversed contrast, with negative values and closest to 0. In T2w images, SSS-fixed brains showed closest values to 0, reflecting less contrast than in the brains fixed with the other two solutions, but still further from 1 than in T1w, translating the overall higher contrast of T2w images.

NBF=Neutral-buffered formalin; SSS=Salt-saturated solution; AFS=Alcohol-formaldehyde solution.

*0.01 < p < 0.05 ; ***p < 0.001 after a Bonferroni correction.

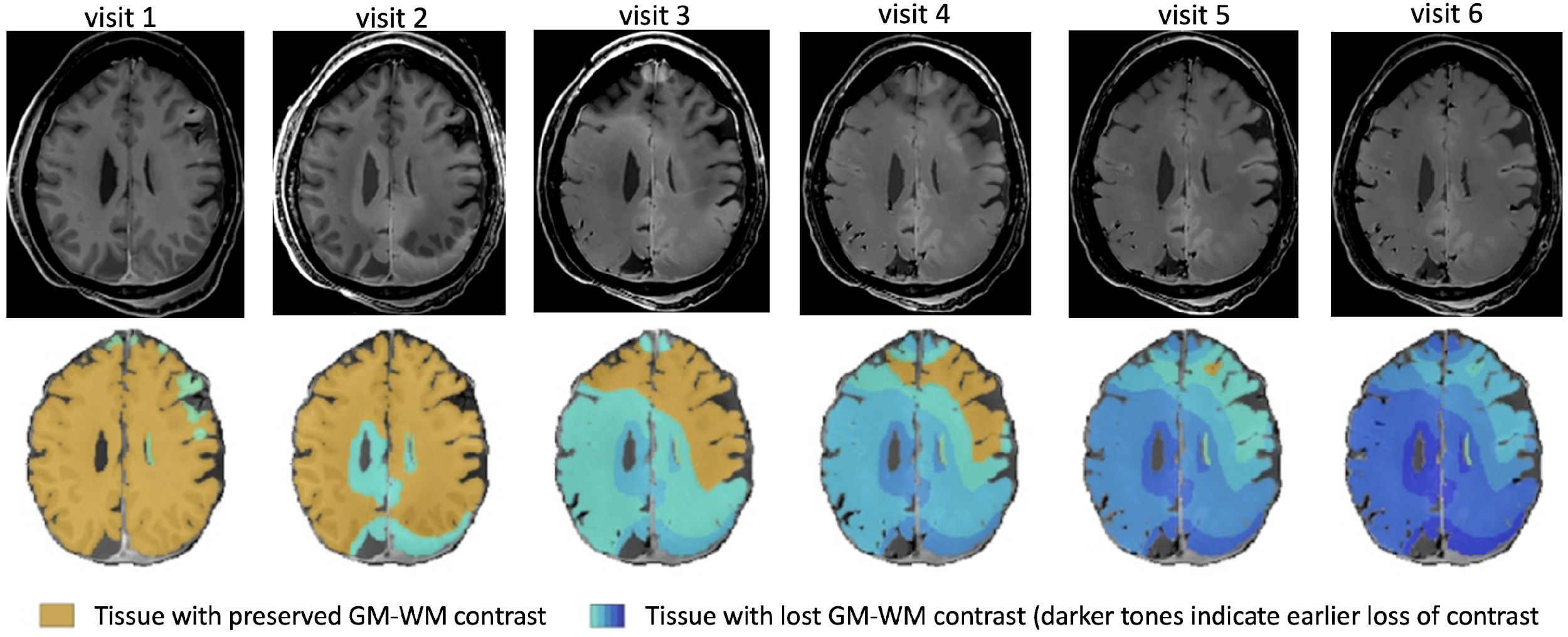

**Supplemental figure 3. Longitudinal assessment of the GM-WM contrast of an *in situ* brain fixed by perfusion with NBF.** This figure was presented at the annual conference of the Society for Neuroscience in 2021 as a virtual poster presentation. First scan (visit 1) was done 2 months after the injection of NBF, which showed no inversion of the contrast (most of the tissue showed preserved GM-WM contrast, reflecting no inversion of the GM-WM). At visit 2, occipital lobes and periventricular spaces showed loss of GM-WM contrast, indicating the beginning of the inversion of the contrast. Inversion of the contrast was completed at visit 6 (8 months after the injection of NBF). (Frigon et al., 2021, Society for Neuroscience).
